# Selection of DNA aptamers targeting PirAB toxin for potential biosensing applications in acute hepatopancreatic necrosis disease (AHPND)

**DOI:** 10.64898/2026.10.14.758549

**Authors:** Vanessa Aguilar-Sánchez, Rylee McKay, Alejandra Hernández-Santoyo, Sonia Araceli Soto-Rodríguez, Jonathan Perreault

## Abstract

The timely detection of AHPND is one of the key factors to avoid the great economic losses that this disease entails for aquaculture. The use of aptamers as detection tools allows us to develop a method focused on the specific needs of the producer, such as being simple and easy to use, and capable of enabling safe, rapid field detection. Specific strains of *Vibrio parahaemolyticus* are the most well-known as the main causal agent of AHPND in shrimp cultures; however, it is not limited to this species, as a transfer of virulence factors has been observed, including the genes encoding the PirAB binary toxin. The concern that this transfer implies has led us to focus our interest on the PirAB binary toxin as a ligand for the development of aptamers capable of identifying it in the field in early stages and avoiding advanced stages of the disease, giving the producer the ability to take preventive actions and reduce the risk of losses.

In the present work we have optimized the SELEX (systematic evolution of ligands by exponential enrichment) aptamer development methodology to rapidly develop candidates capable of detecting both the PirAB complex and its subunits separately, obtaining candidates with a dissociation constant (K_D_) of 65 to 700 nM for the PirB subunit and 1 to 3 μM for the PirAB complex. These aptamers provide the first step toward developing a rapid detection method targeting the causative toxin of AHPND and aid in a new, user-friendly approach to early disease detection.

**Highlights:**

- DNA aptamers were selected against the PirAB toxin of AHPND.
- An accelerated SELEX workflow enabled 17 selection cycles in two weeks.
- Selected aptamers showed preferential binding to PirB (K_D_ 65-700 nM).
- Aptamers targeting PirAB and its subunits were characterized by MST.
- Selected aptamers provide candidates for future PirAB biosensing platforms.

## 1 Introduction

Today, aquaculture is growing more rapidly than all other animal food-production sectors. Its global contribution has increased from 3.9% in 1970 to 33% in 2005 (Ntsama et al., 2018). Due to the high demand for fish products, aquaculture production is increasing, accounting for a larger share of total fish production to ensure supply (Ahmad et al., 2021). Global production of aquatic animals was estimated at 178 million tons in 2020 (FAO, 2020). Specifically, shrimp industry has experienced substantial growth in the last decade; with global imports reaching 4,735,839 tons in 2022, particularly driven by major markets in Asia and the US. Ecuador recorded a compound annual growth rate of 17% from 2012 to 2019 and 25% from 2020 (Villareal, 2023). This expansion, driven by increasing global demand for seafood proteins, has increased interest in its environmental and social impacts and in strategies for more sustainable industry development (González-Rivas & Tapia-Silva, 2023)

Acute hepatopancreatic necrosis disease (AHPND) is the most important bacterial disease of shrimp that has caused mass mortalities in farmed populations of Pacific white shrimp *Penaeus vannamei* and black tiger shrimp *Penaeus monodon* (Tang & Reantaso, 2019; Mai et al., 2021). The causative agents of AHPND are specific strains of the bacteria *Vibrio parahaemolyticus* (abbreviated as Vp_AHPND_) and other *Vibrio* species. The genomes of Vp_AHPND_ contain a 69–73 kb plasmid (known as pVA1) harbouring genes that encode the *Photorhabdus* insect-related (Pir) binary toxin PirABvp. PirAvp (12.7 kDa) and PirBvp (50.1 kDa) are extracellular proteins that act together as a binary toxin. Production losses stemming from outbreaks of AHPND and other shrimp diseases from 2010 to 2016 amounted to approximately US$ 23.6 billion (Tang & Reantaso, 2019). The main strategy for managing AHPND is early diagnosis (Soto-Rodríguez et al., 2015). Histological observations of shrimp affected by AHPND include necrosis, roundness, and massive sloughing of the epithelial cells of the hepatopancreatic tubules into the lumen (pathognomonic lesion), reduction of the reserve vacuoles, and, as the disease progresses, secondary bacterial colonization of the damaged hepatopancreas occurs (Aguilar Rendón et al., 2020). A PCR laboratory test is usually performed before the postlarvae are sent to a farm for cultivation (Soto-Rodríguez et al., 2024). Several PCR protocols for AHPND diagnosis have been developed, most of which are based on the detection of the PirA and PirB genes carried by the pVA-1 plasmid (Peña-Navarro et al., 2020). It is important to note that only diagnostic or research laboratories use this type of method. To date, the most reliable PCR test is the AP4 system, a two-tube nested PCR method targeting the *pirA* and *pirB* toxin genes (Dangtip et al., 2015) but it requires a preincubation step to ensure the detection of the genes (Soto-Rodríguez et al., 2024). A critical component for preventing the spread of AHPND and other shrimp diseases is having available capabilities for timely diagnosis. Early detection of total and pathogenic *Vibrio* is needed and should rely on quick and effective methods (Jusserand et al., 2017).

Molecular methods for detecting PirAB genes can provide high sensitivity and specificity, but their application is limited by the instability and restricted availability of the PirAB region, as well as the dose-dependent toxicity of the PirAB complex (Olvera et al., 2021; González-Gómez et al., 2020; Soto-Rodríguez et al., 2015). Conventional methods, including wet mount analysis, microbiology, and histology, require substantial sample material, equipment, and specialized personnel, while their long processing times can delay intervention and contribute to field losses. Therefore, there is a need for rapid and accessible methods capable of detecting the infection before disease onset or progression.

AHPND develops in four phases: initial, acute, terminal, and remission (Aguilar-Rendón et al., 2020). The initial phase is difficult to detect in the field due to the similarity of the animal’s clinical signs to those of other diseases (WOAH, 2022), which delays the producer’s response until the terminal or chronic phases, thereby considerably increasing mortality and loss rates. Infection is usually diagnosed during its terminal phase, when bacterial loads are elevated as the disease progresses. AHPND is characterized by acute mass mortalities, which usually occur within 30-35 days of stocking grow-out ponds with post-larvae (PL) or juvenile shrimp (WOAH, 2022). AHPND detection is thus critical given the short time farmers have to implement control measures. The chronic (remission) phase has only been observed in some tolerant *Penaeus vannamei* (Aranguren-Caro et al., 2020). By detecting the PirAB binary toxin in shrimp culture samples (i.e., water samples from shrimp ponds) through rapid field monitoring, producers can take appropriate corrective actions.

Aptamers are single-stranded DNA or RNA oligonucleotides selected by Systematic Evolution of Ligands by Exponential Enrichment (SELEX) for their high-affinity and specific binding to target biomolecules (Tuerk & Gold, 1990). Their versatility, relatively low cost, rapid synthesis and target specificity have expanded their applications in biomedical and industrial fields (Chong & Low, 2019), with particular potential in infectious disease management and other niche applications where they may offer advantages over antibodies (Bruno, 2015; Kalita et al., 2023; Ye et al., 2024). DNA-based biosensors and aptamers targeting viruses affecting fish cultures have also demonstrated potential for molecular diagnosis and therapeutic applications (Li et al., 2015). A duplex electrochemical biosensor for the detection of the *tdh* and *trh* genes of *V. parahaemolyticus* in oysters was developed and shows a great sensitivity too, being able to detect as low as 1 CFU of bacteria in oyster homogenate (Campàs et al., 2023). A biosensor using two aptamers and Cut-Assisted rolling circle amplification (CA-RCA) was established for the rapid and visual detection of *V. parahaemolyticus*, enabling detection of as low as 10 CFU/mL or CFU/g in real food samples (Song et al., 2019). Another example of aptamers against *V. parahaemolyticus* was incorporated in the platform of Ahn et al., 2018, this biosensor showed significant activity for detecting and discriminating *V. parahaemolyticus* from other enteric species (Ahn et al., 2018). FAM-labeled aptamer sequences with high binding affinity to *V. parahemolyticus* were identified by flow cytometric analysis, this aptamer displayed a dissociation constant (Kd) of 16.88 ± 1.92 nM, this is another example of an approach that does not require labor-intensive steps (Duan et al., 2012).

These studies demonstrate the potential of nucleic-based strategies for rapid and sensitive detection of *V. parahaemolyticus.* However, detecting the bacterium does not necessarily indicate the presence of the toxins responsible for AHPND. Detecting PirAB could therefore provide information more directly related to disease risk, because AHPND is a toxigenic disease. In addition, approaches requiring PCR and laboratory equipment may limit decentralized applications. Our work explores strategies with potential for simpler and more accessible detection formats. Using SELEX, we identified aptamers that bind the PirAB binary toxin and explored biosensing avenues (Figure 1) for its use as a field detection tool towards the development of a rapid detection method. Additionally, the selection of aptamers has been optimized to be able to perform it in less than two weeks, a much shorter period of time than usual. The candidates selected for selection were characterized to know their affinity and specificity.

**Figure 1.**
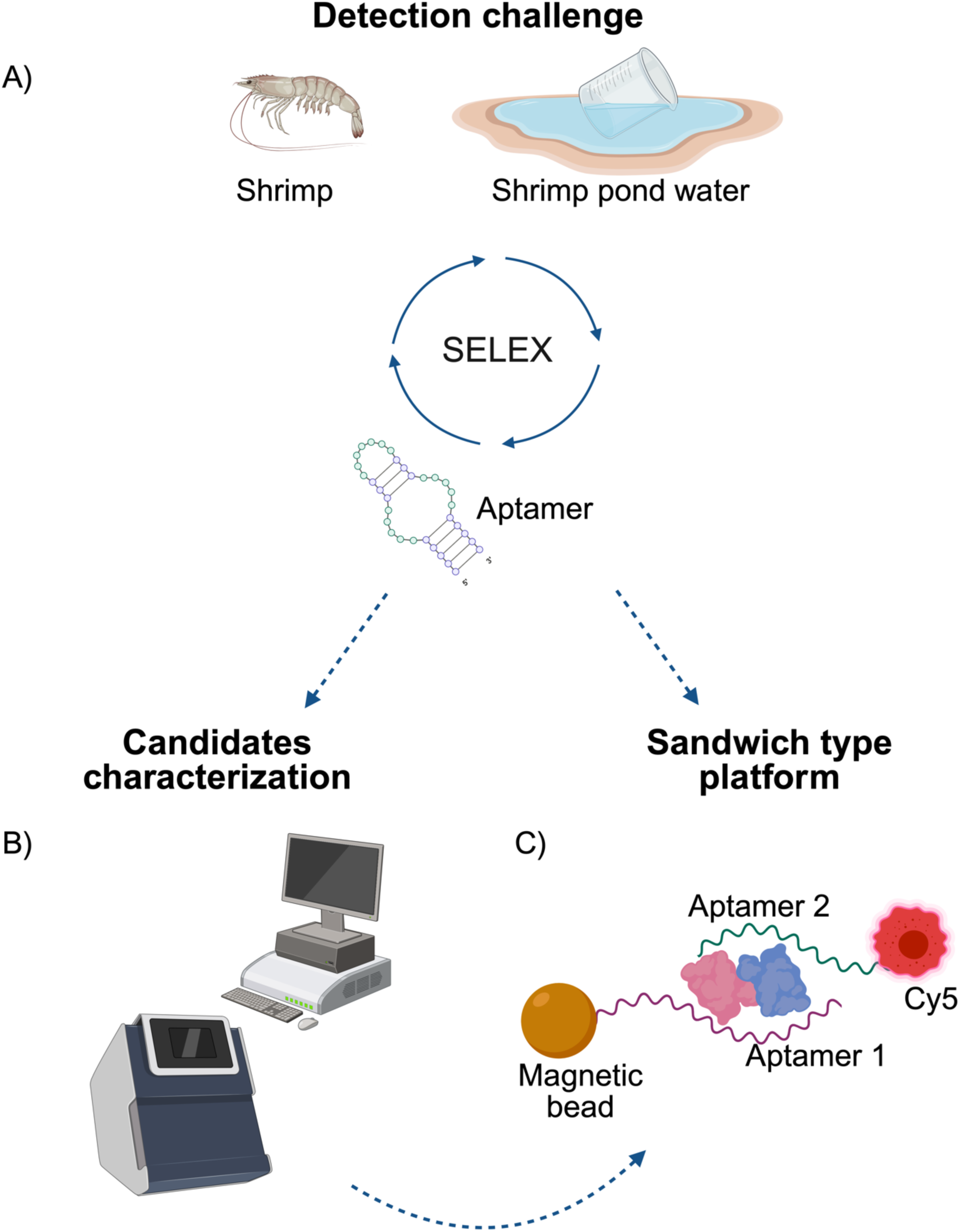
Overview of the strategy. A) Overall SELEX strategy for candidate selection, B) characterization of the candidates by microscale thermophoresis (MST), C) sandwich-type strategy for a platform prototype

## 2 Materials and methods

### 2.1 PirAB toxin

The PirAB complex, PirA and PirB subunits used as targets in SELEX (systematic evolution of ligands by exponential enrichment) methodology were purified in UNAM (Universidad Nacional Autónoma de México), at Chemistry Institute in collaboration with CIAD (Centro de Investigación en Alimentación y Desarrollo) and stocked in sodium phosphate buffer 50 mM, pH 7.5 with 150 mM NaCl.

The strain of *Vibrio parahaemolyticus* M0904 harboring the *pirA* and *pirB* genes (Soto-Rodríguez et al., 2015) was incubated at 30 °C with agitation for 17 hours in soya-tryptone broth (TSB) with 2% NaCl. The cell-free extract was obtained by filtration through a 0.22-µm membrane. Then it was then centrifuged at 10 000 g for 20 min at 4 °C, recovering the supernatant. The supernatant was filtered through a 0.45 μm membrane and applied to a galactose resin (Immobilized D-Galactose resin, Thermo Scientific) previously balanced with sodium phosphate buffer 50 mM (pH 7.5), NaCl 150 mM. Subsequently, the resin was washed with 10 column volumes of the same buffer to remove proteins that did not attach to it. PirAB complex was eluted with galactose 250 mM in the same buffer.

The eluted fraction of galactose resin was dialyzed against a 50 mM sodium phosphate buffer (pH 7.5), 150 mM NaCl, 50 mM galactose and applied to a molecular exclusion column (Agilent Bio SEC-3) balanced with the same buffer where the complex was separated. The purity of the complex was verified by MALDI-TOF mass spectrometry using a matrix of synapinic acid dissolved in a solution of 30% acetonitrile, 70% water, and 0.1% trifluoroacetic acid v/v.

To obtain PirA and PirB separately, the complex was dialyzed against a buffer of K_2_HPO_4_/NaH_2_PO_4_ 0.5 M (pH 7.5) with 1.5 M NaCl. The complex was incubated in this buffer for one night and the fractions separated by ultrafiltration using a 30 kDa-cutoff membrane in an AMICON system. PirA passes through the membrane, while PirB and the complex’s remnants are retained by it. The column was then balanced with sodium-potassium phosphate buffer 200 mM (pH 7.5), NaCl 500 mM.

Both fractions were dialyzed against sodium phosphate buffer 50 mM (pH 7.5), NaCl 150 mM, concentrated by ultrafiltration and the purity of each fraction was verified by MALDI-TOF mass spectrometry using a matrix of synapinic acid dissolved in a solution of 30% acetonitrile, 70% water and 0.1% trifluoroacetic acid v/v.

### 2.2 SELEX library

A 92-nucleotide single-stranded DNA (ssDNA) library containing a 50-nucleotide randomized region (N50) flanked by two constant regions was used as the starting library for SELEX (Figure 2):

**Figure 2.**
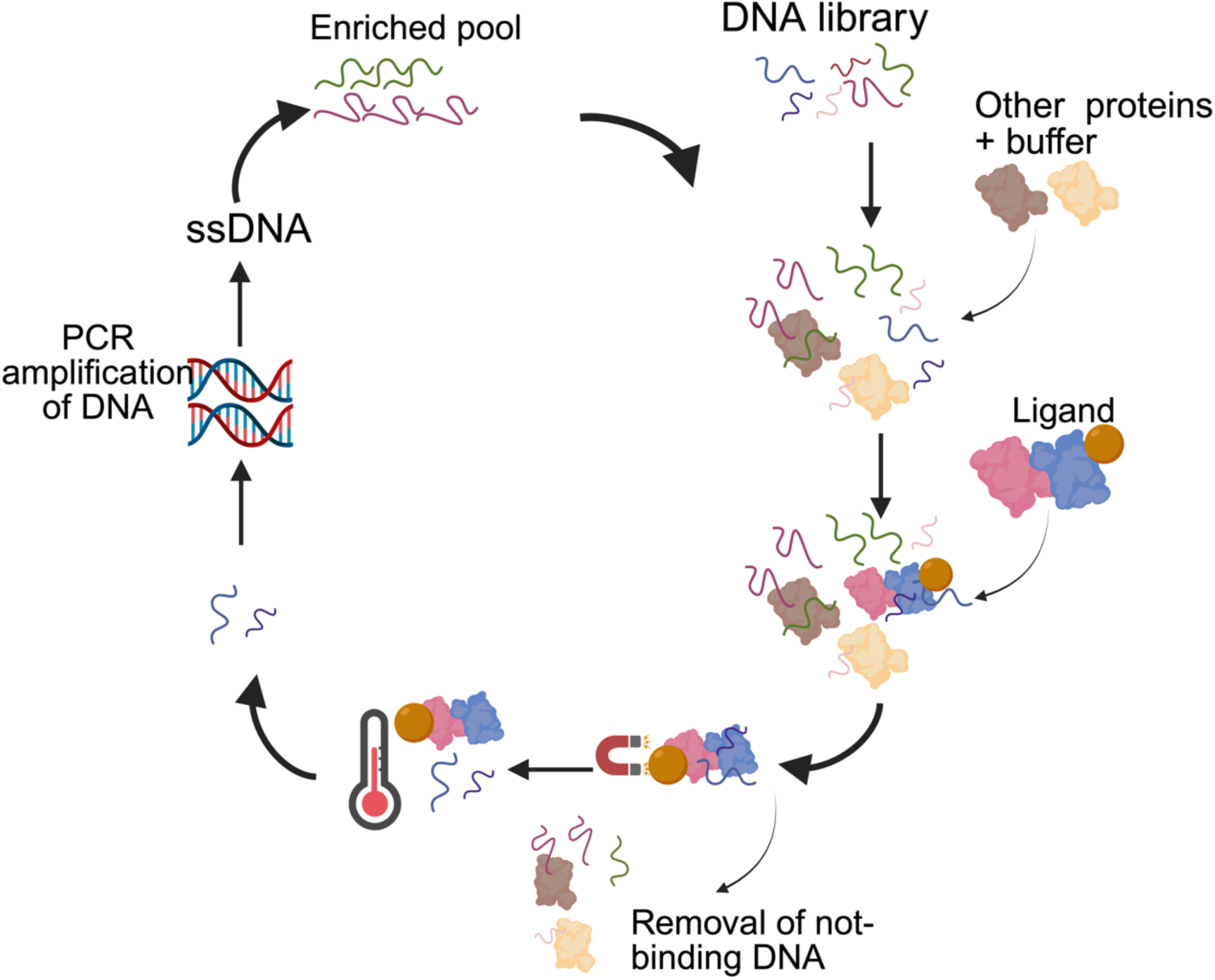
SELEX cycle. A SELEX cycle involves multiple stages, starting with a random library exposed to non-interest targets that titers out candidates that have the ability to bind “non-target molecules” and remove them from enrichment. The remaining sequences are exposed to the ligand of interest and these sequences are eluted from the target and amplified to subsequently generate a single strand DNA and restart the cycle

5’-TTGGTACCTCAGCTATGCAGTG N50 AGAGAGACGGTCCTGCAGAC-3’

The ssDNA library was synthesized by *Integrated DNA Technologies* (IDT). Two primers were used for library amplification: a forward primer (FW) and a 5’-phosphorylated reverse primer (RV):

FW: 5’-TTGGTACCTCAGCTATGCAGTG-3’

RV: 5’-P-GTCTGCAGGACCGTCTCTCT-3’

The 5’ phosphorylation of the reverse primer was introduced to allow selective degradation of the complementary strand by λ-exonuclease following amplification, thereby regenerating the ssDNA library for subsequent SELEX rounds (see Selection section for details).

### 2.3 Target immobilization

PirA, PirB and PirAB (target) immobilization was performed with 200 nm magnetic nanoparticles (NHS modified nanoparticles from Galenvs Sciences): 240 µg of target / mg of magnetic nanoparticles (MNPs) was mixed) in 500 µl of coupling buffer (0.1 M NaHCO_3_, 0.5 M NaCl pH 8.3) and incubated with rotation at room temperature 1 hour. Following this incubation, MNPs were immobilized on the side of the tube by a magnet and washed two times with 1 ml of coupling buffer and blocked with 1 ml of 0.1 M Tris/HCl pH 8.0 with incubation in rotation at room temperature 1 hour. The sample was further washed with 1 ml of 0.1 M Tris/HCl (pH 8.0) and 1 M NaCl, and after with 1 ml of acetate buffer (0.1 M glacial acetic acid and 1 M NaCl, pH 4.0) and finally with 1 ml of working buffer (20 mM Tris/HCl, 100 mM NaCl, 5 mM KCl, 2 mM CaCl_2_ and 1 mM MgCl_2_ (pH 7.4)). The final suspension was prepared in 300 µl of working buffer, resulting in a target concentration of 14.6 µM target bound to MNPs, based on an estimated binding efficiency of 60%, for use in positive selection rounds.

### 2.4 Selection

A total of 17 SELEX cycles were performed with the conditions presented in Table 1.

**Table 1.** SELEX cycles.

| Cycle | Elution | Incubation time (min) | Positive selection (target volume, µl) | Negative selection (µM) |
| --- | --- | --- | --- | --- |
| Series 1: PirAB complex |  |  |  |  |
| 1 | - | 60 | 200 | - |
| 2 | - | 60 | 100 | - |
| 3 | - | 60 | 50 | - |
| 4 | - | 45 | 20 | 0.25 BSA + lysozyme |
| 5 | - | 45 | 20 | 0.25 BSA + lysozyme |
| 6 | + | 30 | 20 | 0.25 BSA + lysozyme |
| 7 | + | 30 | 20 | 0.25 BSA + lysozyme |
| 8 | + | 30 | 20 | 0.25 BSA + lysozyme |
| 9 | + | 20 | 20 | 0.25 BSA + lysozyme |
| 10 | + | 20 | 15 | 0.5 lysozyme |
| 11 | + | 20 | 15 | 0.5 lysozyme |
| 12 | + | 15 | 10 | 0.5 lysozyme |
| 13 | + | 15 | 10 | 0.5 lysozyme |
| 14 | + | 10 | 5 | 0.5 lysozyme |
| Series 2: Subunit PirA |  |  |  |  |
| 15 | + | 10 | 5 | 0.25 BSA + lysozyme |
| 16 | + | 10 | 5 | 0.25 BSA + lysozyme |
| Series 2: Subunit PirB |  |  |  |  |
| 15 | + | 10 | 5 | 0.25 BSA + lysozyme |
| 16 | + | 10 | 5 | 0.25 BSA + lysozyme |
| Series 3: PirAB complex |  |  |  |  |
| 17 | + | 10 | 5 | 0.25 BSA + lysozyme |
| Target volume: number of μl taken from a 5% MNP suspension. | | | | |

Positive selections were made with the volumes of immobilized target suspensions and the different incubation times at room temperature showed in Table 1. After positive selection, an elution (started in SELEX cycle #6) was done for 5 minutes at 94 °C to separate the candidates from nanoparticles.

Negative selection was performed using BSA (Bovine Serum Albumin) and lysozyme in solution in working buffer at different concentrations (details are provided in Table 1).

After each selection, the double stranded DNA production was done by PCR with the following final concentrations: 0.5 μM of each primer, 200 μM of each dNTP (New England BioLabs, NEB), 1 X of PCR buffer (NEB), 1 U of Accustart II Taq DNA Polymerase (NEB) and 2 μl of DNA template (SELEX library) in a final volume of 50 μl. The amplification conditions used were initial denaturation at 94 °C for 3 minutes, denaturation at 94 °C for 30 seconds, hybridization at 58 °C for 30 seconds, elongation at 72 °C for 30 seconds and a final elongation at 72 °C for 3 minutes for 25 PCR cycles.

The single stranded DNA production was done with λ-exonuclease (NEB) digestion, using 3 μl of enzyme (at 5000 U/ml) per 50 μl of PCR product, incubating 30 minutes at 37 °C, followed by an inactivation of 10 minutes at 80 °C to immediately start the new cycle.

### 2.5 Sequencing

Following SELEX, generations 6, 10 and 14 (from series 1: PirAB), 17 (from series 3: PirAB), 16 (from series 2: PirA) and 16 (from series 2: PirB) were sequenced (CERMO-FC services at UQAM and Innovation Centre Genome Québec, Montreal, Canada) and the results were analyzed by the Aptasuite software (Hoinka et al., 2018), aligning the reads with the primers used during the selection of the library by the software and obtaining as a result the following data: number of reads, enrichment in each cycle and mutations in the random sequence, and others.

### 2.6 Microscale thermophoresis

The selected candidate APT4 was synthetized and modified with Cy5 (Alpha DNA, Montreal, Canada), and other selected candidates; A458, B89, AB115 and AB171 were modified with Cy5 in the laboratory by PCR previously described in 2.4 Selection section with the following primers prepared by *Integrated DNA Technologies* (IDT): FW: 5’-Cy5-TTGGTACCTCAGCTATGCAGTG; RV: 5’-P-GTCTGCAGGACCGTCTCTCT. Followed by a λ-exonuclease digestion as previously described to generate a Cy5-labeled single-strand DNA corresponding to the selected aptamers.

An aptamer concentration of ∼5 nM was incubated in a serial dilution of its target: PirAB complex (7 μM to 0.2 nM), PirA subunit (12 μM to 0.5 nM) or PirB subunit (65 μM to 2 nM) in optimized working buffer (PBS 1x containing MgCl_2_ 5 mM) in rotation at room temperature for 20 minutes, following an incubation of 24 hours at 4 °C for a correct reading by Monolith NT.115 Pico device (NanoTemper technologies) to analyze their interaction. The device generates a *K*_D_ value based on variation in fluorescence and variations in diffusion upon sample heating.

## 3 Results

### 3.1 Selection of aptamers against PirAB

Aptamers were selected using the SELEX methodology (Figure 2) against the binary toxin PirAB, purified from the culture of the *Vibrio parahaemolyticus* M0904 strain. To accelerate the selection process, ssDNA was generated by λ-exonuclease digestion of PCR products obtained using 5’-phosphorylated reverse primer, enabling selective degradation of the phosphorylated complementary DNA strand and avoiding the conventional PAGE purification step. This streamlined workflow allowed us to perform two or three SELEX rounds per day, depending on the selection time. Five sequences were chosen as candidates Series 1, and four sequences Series 2 and Series 3, based on an Aptasuite analysis (0.9.8 version) (Hoinka et al., 2018) (Table 2). A preliminary characterization for the candidates selected from SELEX Series 1 was performed using a sandwich test (details of this assay can be found in the Supplementary material), we have selected two candidates from series 1 (APT3 and APT4) that could work together to identify the toxin complex. Candidate APT4 was selected based on the binding percentages vs PirAB complex and vs magnetic beads as negative controls (Supplementary Table S3). The complete sequences of the selected aptamers are shown in table 2.

**Table 2.**
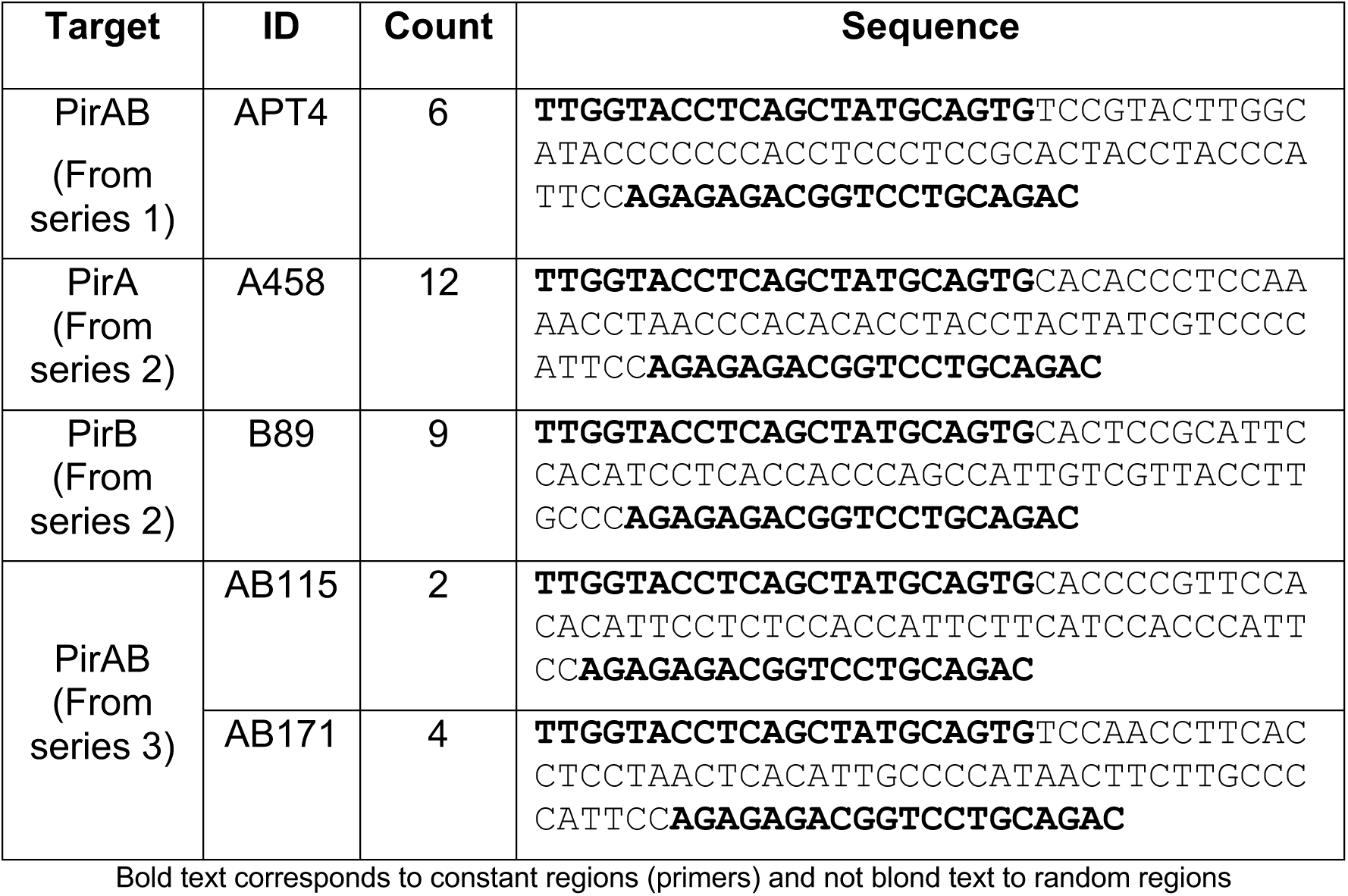
PirAB complex, PirA subunit and PirB subunit candidates.

In principle, the combined use of two aptamers that each bind different epitopes of the PirAB complex would permit a sandwich assay (as schematized in Figure 1C). This can potentially be performed by combining aptamers that bind different epitopes of the same subunit; or by combining aptamers against each subunit of the toxin. Overall, because the epitopes recognized by the candidate aptamers are not known, many combinations were evaluated, but none of the 20 tested combinations (Supplementary Table S2) gave conclusive results (Supplementary Table S1).

Because the combined use of these different aptamers had limited success in sandwich assays, we performed additional SELEX cycles (Figure 3) to isolate aptamers against PirA and PirB separately (Series 2: PirA and Series 2: PirB). The rationale being that combining an aptamer against subunit A and another one against subunit B would more likely yield a successful sandwich assay. For good measures, we also added cycle 17 (with a mix of PirA and PirB libraries) with the complex PirAB again to ensure compatibility with the complete toxin (Series 3), i.e., that the targeted epitopes remain accessible within the complex. The newly selected candidates were initially characterized by microscale thermophoresis (MST) to identify candidate pairs with complementary binding proprieties that could be further evaluated in future work to improve the sandwich assay and potentially adapted to a biosensing platform.

**Figure 3.**
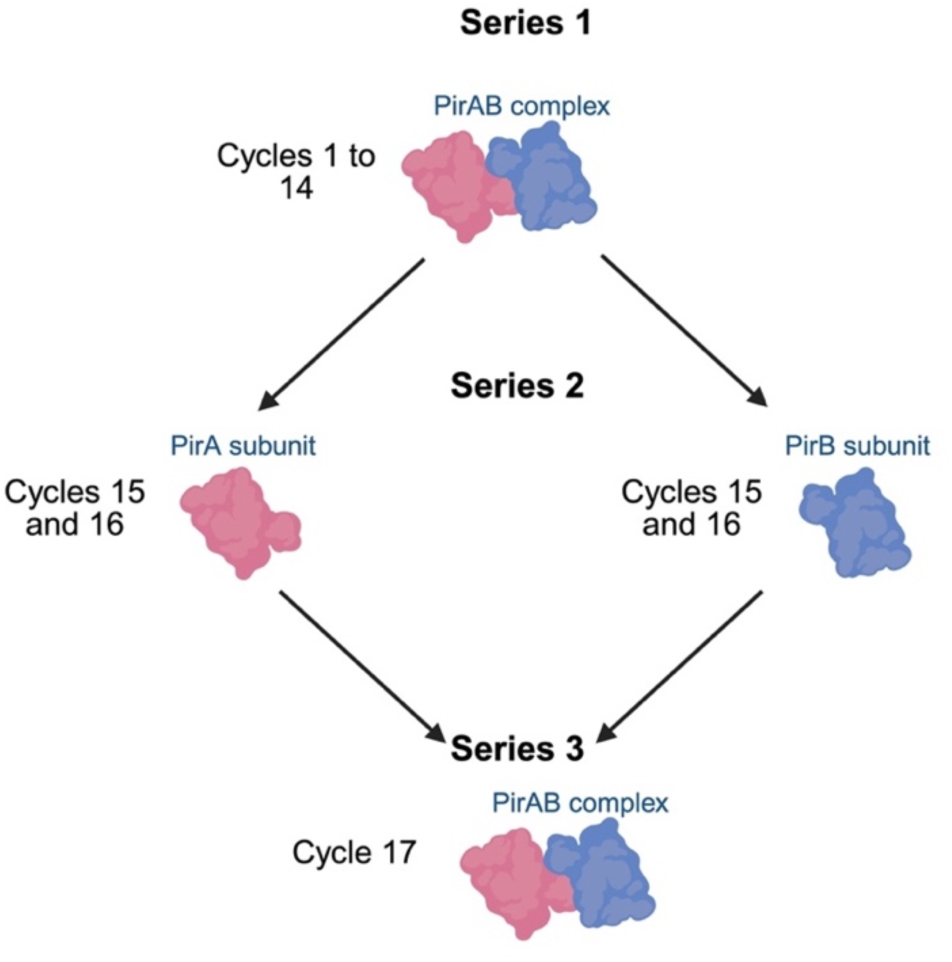
SELEX series performed. Target used in the different SELEX cycles performed

### 3.2 Candidate aptamers characterization by microscale thermophoresis

Selected candidates: APT4, A458, B89, AB115 and AB171 (Table 2), were characterized against the PirAB complex and against the PirA and PirB subunits to evaluate the affinity of each candidate. The selected aptamers have affinities in the nanomolar range (up to low micromolar) for the toxin and its specificity was verified using different negative controls. Table 4 shows a summary of K_D_ results for the target and for bovine serum albumin (BSA) as the more representative negative control. B89 and APT4 show the ability to identify PirAB complex, while B89, AB115 and AB171 have shown a more interesting *K*_D_ against the PirB subunit (Figure 4).

**Table 4.** Dissociation constants, signal-to-noise ratios, and response amplitudes of selected aptamers against PirAB toxin, its subunits, and BSA as a negative control.

| SELEX<br>series<br>origin | | PirA | | | PirB (nM) | | | PirAB (μM) | | | BSA (μM) | | |
| --- | --- | --- | --- | --- | --- | --- | --- | --- | --- | --- | --- | --- | --- |
| | | $K_D$<br>(μM) | S/N | RA | $K_D$<br>(nM) | S/N | RA | $K_D$<br>(μM) | S/N | RA | $K_D$<br>(μM) | S/N | RA |
| Series<br>1 | **APT4** | 9.88 | 41 | 71 | -- | -- | -- | 1.27 | 18 | 20 | -- | -- | -- |
| Series<br>2 | A458 | 5.21 | 42 | 43 | 698 | 15 | 51 | -- | -- | -- | 3.9 | 5 | 4 |
|  | **B89** | 6.92 | 50 | 61 | 64.3 | 13 | 31 | 3.13 | 9 | 14 | 11.6 | 8 | 5 |
| Series<br>3 | AB115 | 6.27 | 71 | 65 | 182 | 16 | 48 | -- | -- | -- | 1.7 | 7 | 6 |
|  | AB171 | 7.64 | 8 | 85 | 181 | 12 | 74 | -- | -- | -- | -- | -- | -- |

Summary of the dissociation constants ( $K_D$ ), signal-to-noise ratios (S/N), and response amplitudes (RA) of the selected aptamers against the target proteins and bovine serum albumin (BSA) as an example negative control showing binding, the rest of negative controls without binding (lysozyme, hemicellulase, heparin, insulin, dextran sulfate, Tris-HCl and polyhistidine-tag) are provided in the Supplementary Material.

**Figure 4.**
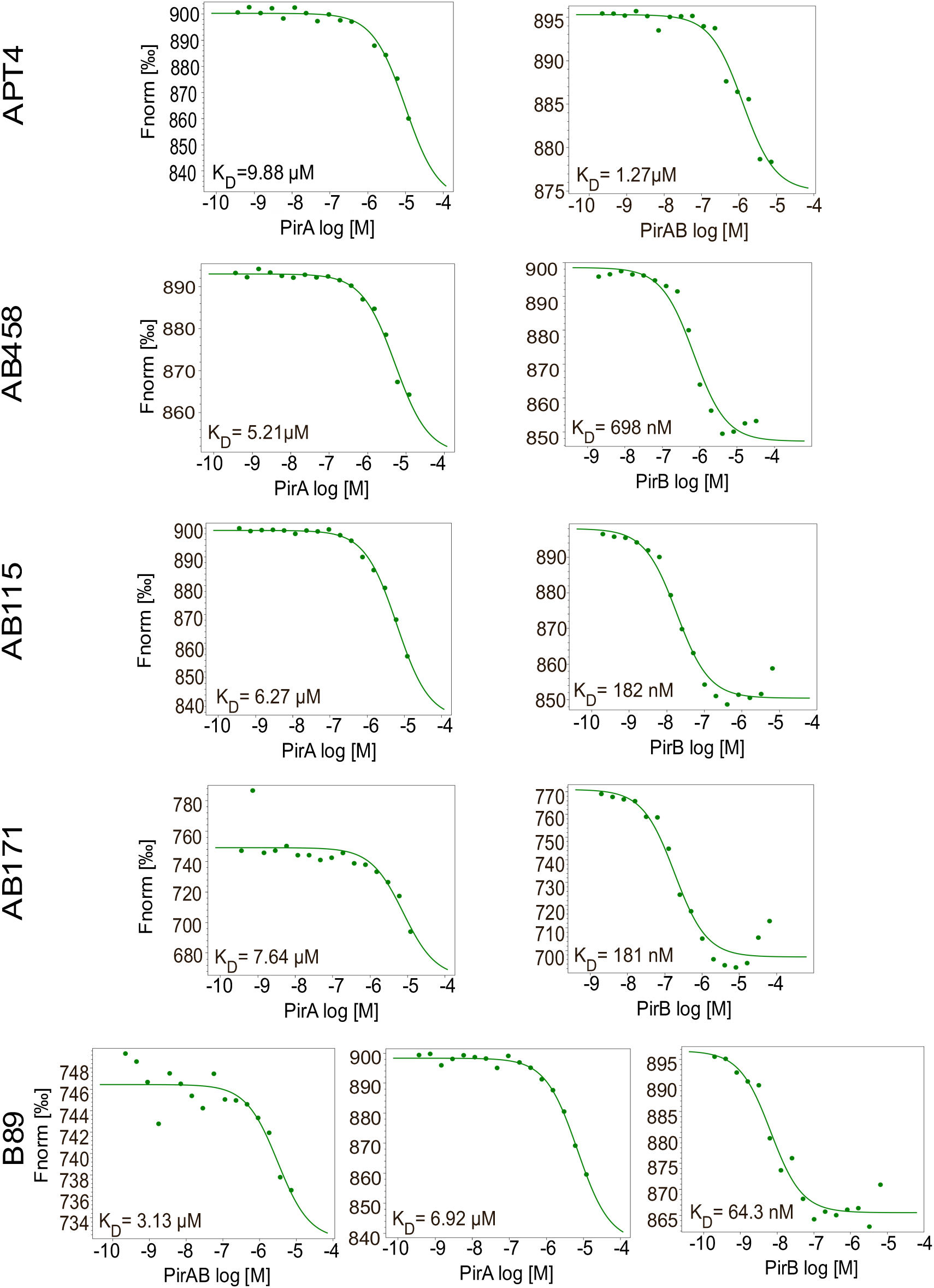
Affinity of selected candidates. Binding affinity of selected candidates: each candidate was assayed by microscale thermophoresis in presence of PirAB complex, PirA subunit or PirB subunit.

The curves against several proteins used as negative controls: BSA, lysozyme, hemicellulase, heparin and insulin, dextran sulfate as well as against Tris-HCl (used during blocking step of target immobilization), and TEV protease protein in presence of polyhistidine-tag (which was present in our ligand during candidate selection) are available in supplementary material (Supplementary Figure S2). These were used in order to verify the specificity of the aptamers for PirAB and subunits. It was not possible to determine *K*_D_ values for most of these “negative control proteins” (or the signal to noise ratio was not adequate), confirming specificity against the toxin and subunits.

## 4 Discussion

The methodology used in the present work for aptamer enrichment allowed us to perform 17 selection cycles within a two-week period. This accelerated workflow was achieved by combining magnetic bead-based target separation with direct enzymatic generation of ssDNA. In particular λ-exonuclease digestion of PCR products generated with 5’-phosphorylated reverse primer eliminated the need for gel-based ssDNA preparation, reducing the time and handling required between successive selection cycles. This approach has previously been described as an efficient method for ssDNA generation (Hao et al., 2020); while magnetic bead-based SELEX strategies have also been reported to facilitate rapid target-bound and unbound fraction separation (Hünniger et al., 2014); (Ayass et al., 2023).

In our workflow, this strategy also provided a rapid and cost-effective approach for generating Cy5-labeled ssDNA for microscale thermophoresis (MST) assays, avoiding the need to order individually labeled aptamers. Although the resulting ssDNA preparations may contain residual reaction components or incompletely processed material, this approach enabled the rapid characterization of newly generated candidates and facilitated the iterative screening of aptamer–target interactions during the selection process.

Characterization of the selected candidates demonstrated measurable binding to our ligand and revealed a clear preference for PirB, with apparent dissociation constants ranging from 65 to 700 nM, compared with approximately 3 µM for PirAB. Some candidates also showed measurable binding to PirA, with apparent dissociation constants ranging from 6 to 10 µM, indicating weaker recognition of this subunit compared with PirB. The preferential recognition of PirB may be partly related to its larger size compared with PirA (50.1 kDa and 12.7 kDa, respectively). A larger protein subunit may offer a greater molecular surface and wider range of potential interaction sites for aptamer binding, although protein size alone does not determine binding affinity. Differences in the surfaces accessibility, local conformation, and distribution of charged or polar residues between the two subunits may also influence the interactions established with the selected aptamers. In addition, the heterotetrameric structure of PirAB complex (Soto-Rodríguez et al., 2022) may contribute to the observed binding profile, as its multimeric organization could potentially mask or reduce the accessibility of PirA while leaving PirB more exposed to aptamer binding. Furthermore, the conformation adopted by the assembled complex may alter the accessibility or presentation of epitopes recognized by aptamers compared with those available on isolated subunits. This could help explain why some candidates displayed stronger apparent affinity for PirB than for the PirAB complex. This structural arrangement could therefore partly explain the preferential recognition of PirB observed among the selected candidates.

During the SELEX process, substantial library losses were observed at certain selection cycles. This may be partly related to the absence of purification steps, as the workflow was intentionally explored with the aim of developing a more rapid SELEX procedure. This may also help explain why some aptamers, particularly those from series 3, did not show measurable affinity for PirAB complex despite having undergone several selection cycles against this target before subsequent selection toward individual units. In addition, sequencing conditions were not optimal, resulting in a relatively limited number of reads, with only a few hundred thousand reads obtained. This reduced sequencing depth limited the assessment of sequence enrichment and may have affected the identification of representative candidates.

Of the candidates characterized by MST, only APT4 was evaluated in the sandwich assay, which did not result in a functional assay under the conditions tested, while the remaining candidates were selected subsequently and have not yet been evaluated in this format. The binding profiles of the candidates nevertheless provide a basis for further optimization of aptamer performance and exploration of their potential incorporation into future biosensing platforms. The moderate affinities observed may currently limit their direct application for highly sensitive toxin detection in field samples; however, further optimization of aptamer selection and assay conditions such as seawater and shrimp tissue could improve their performance for PirAB-related detection.

Importantly, the binding experiments also indicate that the candidates exhibit a preferential interaction with the selected target, even when some degree of binding to other tested proteins was observed. Rather, the relative differences observed in the affinity curves suggest that the selected aptamers recognize PirAB or its subunits more strongly than the non-target proteins evaluated. This preferential binding is particularly relevant for the development of detection platforms intended for complex samples, where the complete absence of nonspecific interactions is difficult to achieve. If this preferential recognition is maintained in relevant matrices, the aptamers could potentially discriminate the target from other abundant proteins and enable detection of low concentrations of PirAB/subunits. However, this hypothesis will require further validation through systematic selectivity studies, matrix-effect assessments and determination of the analytical sensitivity of the resulting assay.

Aptamers offer several advantages as molecular recognition elements for biosensing applications, including in vitro selection, chemical synthesis, sequence modification, and compatibility with different signal-transduction platforms. In this study, the rapid generation of Cy5-labeled ssDNA and the characterization of candidates with preferential binding to PirB provide a foundation for further development of aptamer-based detection strategies for AHPND-associated toxins. Although additional optimization will be required to improve binding performance and stablish analytical sensitivity and selectivity in complex matrices, these results support the continued exploration of the selected aptamers as potential recognition elements for PirAB-related biosensing applications.

## 5 Conclusions

The present results therefore support the continued investigation of these aptamers as recognition elements for rapid PirAB toxin detection. Further optimization of aptamer affinity, target selectivity, assay configuration and signal amplification will be necessary to determine whether these candidates can achieve the sensitivity and robustness required for field applications. Nevertheless, the results obtained here indicate that the selected aptamers have sufficient target recognition to justify their continued evaluation in prototype biosensing platforms.

A rapid detection method could contribute to earlier identification of AHPND-associated toxin and, consequently, provide shrimp farmers with information that could support timely management decisions during periods of disease risk. Depending on the production system and applicable management practices, such information could facilitate interventions such as adjustments to feeding strategies, use of appropriate water-quality management measures, or decision regarding harvest timing. The value of such interventions would ultimately depend on the capability of the platform to provide sufficiently rapid reliable information.

In aquaculture, reported aptamer-based detection approaches for *Vibrio parahaemolyticus* have primarily focused on the identification of the bacterial pathogen rather than on the detection of PirAB toxin. However, the presence of *V. parahaemolyticus*, or even of the *pirA* and *pirB* genes, does not necessarily indicate the secretion of the toxin responsible of AHPND. In contrast, the present work focuses on the recognition of the PirAB toxin and its subunits at the protein level, providing a complementary approach for assessing the presence of the virulence factor directly associated with AHPND.

Several nutritional and microbial-based strategies have been investigated or commercially implemented to support shrimp health. For example, the fermented corn protein concentrate (FCPC) commercially known as MOTIV® has been reported to decrease bacteria loads in the hepatopancreas of shrimp *Litopenaeus vannamei* and to improve the survival under salinity stress, together with effects on antibacterial response (Novriadi et al., 2022). Probiotics have also been investigated as alternatives or complementary approaches to antimicrobial treatments in aquaculture, with reported effects on shrimp health and disease resistance (Madhana et al., 2021). Other examples include *Bacillus subtilis* E20 or *Bacillus albus* SMG-1 and *Bacillus safensis* SMG-2 supplementation to improve growth and nutrient utilization in shrimp (Tsai et al., 201; Kim et al., 2021). More specifically, certain probiotics have been evaluated against AHPND-causing *Vibrio parahaemolyticus* strains. For example, dietary supplementation with *Bacillus* spp. and the use of host-derived lactic acid bacteria have been reported to improve shrimp immune responses and resistance to AHPND-associated bacterial challenge (Kewcharoen, 2019; Chomwong, 2018). In addition, the antimicrobial compound amicoumacin A, produced by *Bacillus subtilis* BSXE-1601, has shown inhibitory activity against shrimp-pathogenic vibrios, including an AHPND-causing strain (Wang et al., 2020).

The implementation of such preventive and management strategies could therefore benefit from rapid diagnostic tools capable of providing actionable information on the sanitary status of shrimp cultures. In this context, although additional optimization and validation are required, the aptamers identified in the present study represent a starting point for the development of portable and cost-effective biosensing platforms.

## Credit authorship contribution statement

**Vanessa Aguilar-Sánchez**: Conceptualization, Investigation, Methodology, Validation, Visualization, Writing – original draft. **Rylee McKay**: Investigation, Validation, **Alejandra Hernández-Santoyo**: Resources. **Sonia Araceli Soto-Rodríguez**: Resources, Writing – Review & Editing. **Jonathan Perreault**: Conceptualization, Funding acquisition, Supervision, Writing – Review & Editing.

## Acknowledgements

The authors acknowledge the support of the Instituto de Química, Universidad Nacional Autónoma de México (UNAM), and the Centro de Investigación en Alimentación y Desarrollo (CIAD), Mexico, in connection with the preparation of the PirAB toxin complex and its PirA and PirB subunits used in this study.

This work was supported by the Natural Sciences and Engineering Research Council of Canada (NSERC) (grant number ALLRP 590410) and PRIMA Québec (project number R25-13-004-PME). VAS received support from the Globalink Graduate Fellowship and the Armand-Frappier Foundation Graduate Scholarship, while RM received support through an NSERC undergraduate summer research fellowship.

