## Supplementary Material for "Selection of DNA aptamers targeting PirAB toxin for potential biosensing applications in acute hepatopancreatic necrosis disease (AHPND)"

---

#### Sandwich test as preliminary selection

One version (N) of five selected aptamer candidates (Table S1) was modified with a $NH_2$ group in order to bind it to MNPs. Another version (C) of the same five selected aptamer candidates was labelled with Cy5 to have a fluorescence signal once the target has been detected, to have an overall scheme: magnetic bead-$NH_2$ aptamer-PirAB complex-aptamer Cy5 (Figure S1). We have measured the fluorescence signal with a Typhoon FLA9500 and quantified it by ImageJ.

$NH_2$ aptamer candidate versions were immobilized using 3% NHS beads (magnetic beads that allows covalent amine-based conjugation) and 50 pmoles of candidate following the target immobilization protocol described above. Then, a PirAB complex concentration of 0.3 mg/ml was added to the mix and incubated for 15 minutes, in rotation at room temperature. Once this initial binding was done, we added 5 pmol of Cy5-aptamer candidate to 5 µl of the MNP-aptamer-PirAB complex. Table 3 describes the evaluated combinations (a total of 20 samples were tested to verify the joint work of each aptamer candidate) (Table S2).

According to the results in the sandwich tests, the aptamer candidates 3N and 4C present a better response in the joint work of toxin identification (Table S2), the results of the tests with both candidates together were similar and with the highest percentages of binding. Given these results we have chosen both candidates as the most appropriate to continue the characterization work. In addition, aptamer 4 presents one of the lowest percentages in the negative control tests against magnetic beads, it was selected for microscale thermophoresis characterization.

**Table S1** PirAB complex candidates.

| Target | ID | Count | Sequence |
|---|---|---|---|
| PirAB (From series 1) | APT1 | 30 | **TTGGTACCTCAGCTATGCAGTG**TAGAGAGACGGTCCTGCAGAC**AGAGAGACGGTCCTGCAGAC** |
| | APT2 | 6 | **TTGGTACCTCAGCTATGCAGTG**CCCCCCATCCTCCTTACAGCGCCCCCCACATCCTTACCACCCCATTCC**AGAGAGACGGTCCTGCAGAC** |
| | APT3 | 6 | **TTGGTACCTCAGCTATGCAGTG**GGGGGAAGTACTTAGTGTGTTGTGGT**AGAGAGACGGTCCTGCAGAC** |

| | | | |
|---|---|---|---|
| | APT4 | 6 | **TTGGTACCTCAGCTATGCAGTG**TCCGTACTTGGCATACCCCCCCACCTCCCTCCGCACTACCTACCCATTCC**AGAGAGACGGTCCTGCAGAC** |
| | APT5 | 6 | **TTGGTACCTCAGCTATGCAGTG**TGGGACAAGGAAGTAATGTGTGTTGTGGTA**AGAGAGACGGTCCTGCAGAC** |

Bold text corresponds to constant regions (primers) and not blond text to random regions

**Table S2** Sandwich test relation

| Aptamer version | Sandwich test combinations | | | | |
|---|---|---|---|---|---|
| N | APT1 | APT2 | APT3 | APT4 | APT5 |
| C | APTs: 2, 3, 4, 5 | APTs: 1, 3, 4, 5 | APTs: 1, 2, 4, 5 | APTs: 1, 2, 3, 5 | APTs: 1, 2, 3, 4 |

(For detailed sequences look at Table 2).

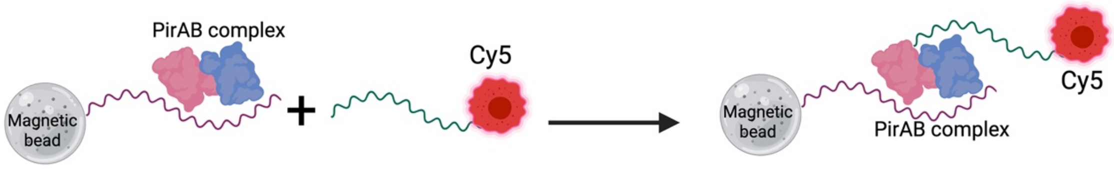

**Figure S1** Sandwich test principle

Table S3 Sandwich test results

| Mix Aptamer | Bound aptamer (%) |
|---|---|
| 1N + 2C | 6 |
| 1N + 3C | 13 |
| 1N + 4C | 6 |
| 1N + 5C | 0 |

| Mix Aptamer | Bound aptamer (%) |
|---|---|
| 2N + 1C | 7 |
| 2N + 3C | 3 |
| 2N + 4C | 6 |
| 2N + 5C | 2 |

| Mix Aptamer | Bound aptamer (%) |
|---|---|
| 3N + 1C | 5 |
| 3N + 2C | 3 |
| 3N + 4C | 6 |
| 3N + 5C | 2 |

| Mix Aptamer | Bound aptamer (%) |
|---|---|
| 4N + 1C | 5 |
| 4N + 2C | 6 |
| 4N + 3C | 3 |
| 4N + 5C | 5 |

| Mix Aptamer | Bound aptamer (%) |
|---|---|
| 5N + 1C | 5 |
| 5N + 2C | 2 |
| 5N + 3C | 3 |
| 5N + 4C | 4 |

| Binding to magnetic beads (C-) | |
|---|---|
| Aptamer | Bound aptamer (%) |
| 1 | 1 |
| 2 | 6 |
| 3 | 2 |
| 4 | 1 |
| 5 | 2 |

xN/xC: x: candidate number, N:$NH_2$ aptamer version; C:Cy5 aptamer version.

$NH_2$ version conjugated to magnetic beads and PirAB complex tested with all other Cy5 version candidate aptamers;

C-: Negative control; all Cy5 version candidate aptamers tested vs magnetic beads.

*Note that all tests were done in duplicate, so no error bars are shown.

### Specificity test

**Figure S2** Specificity test

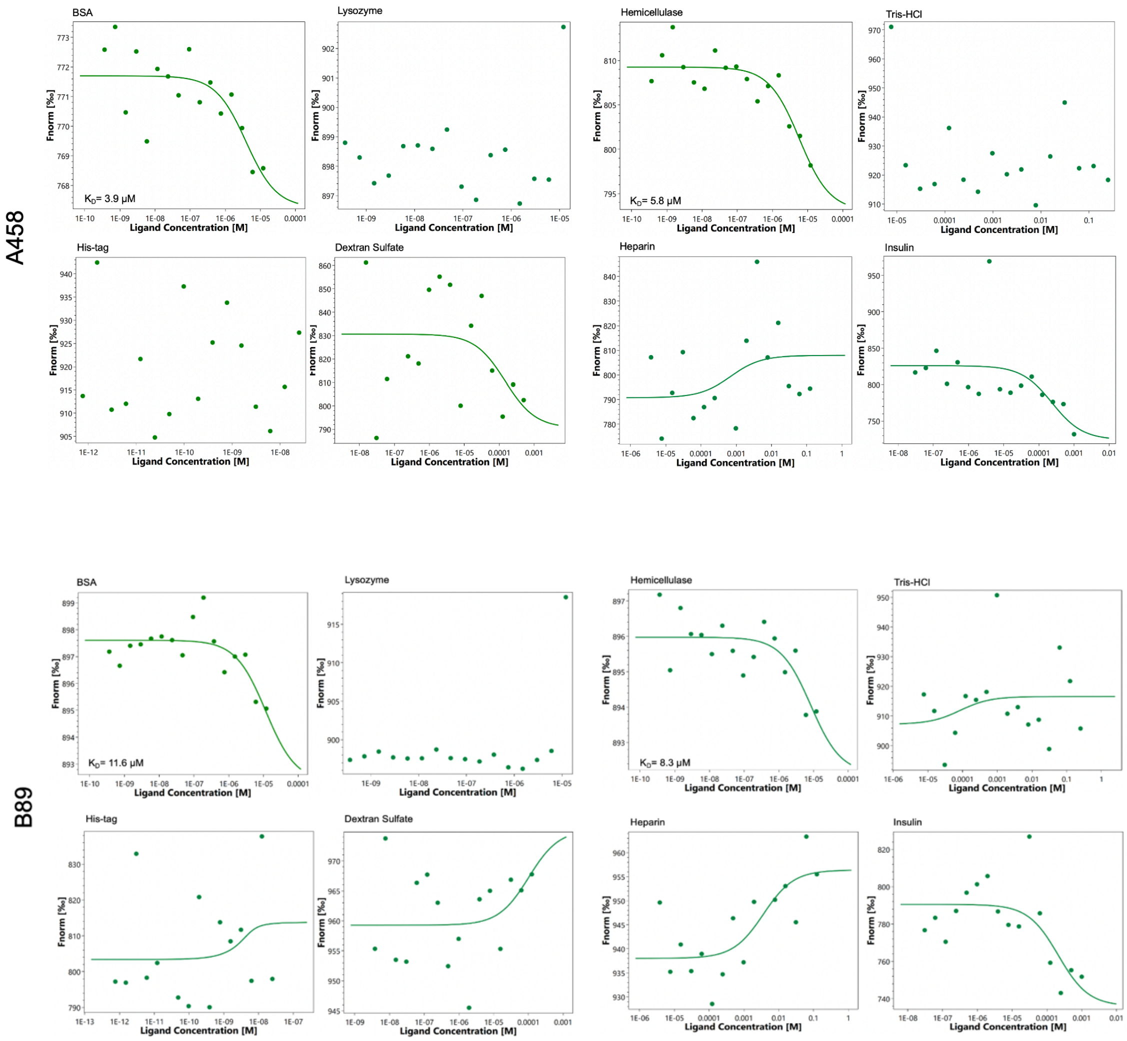

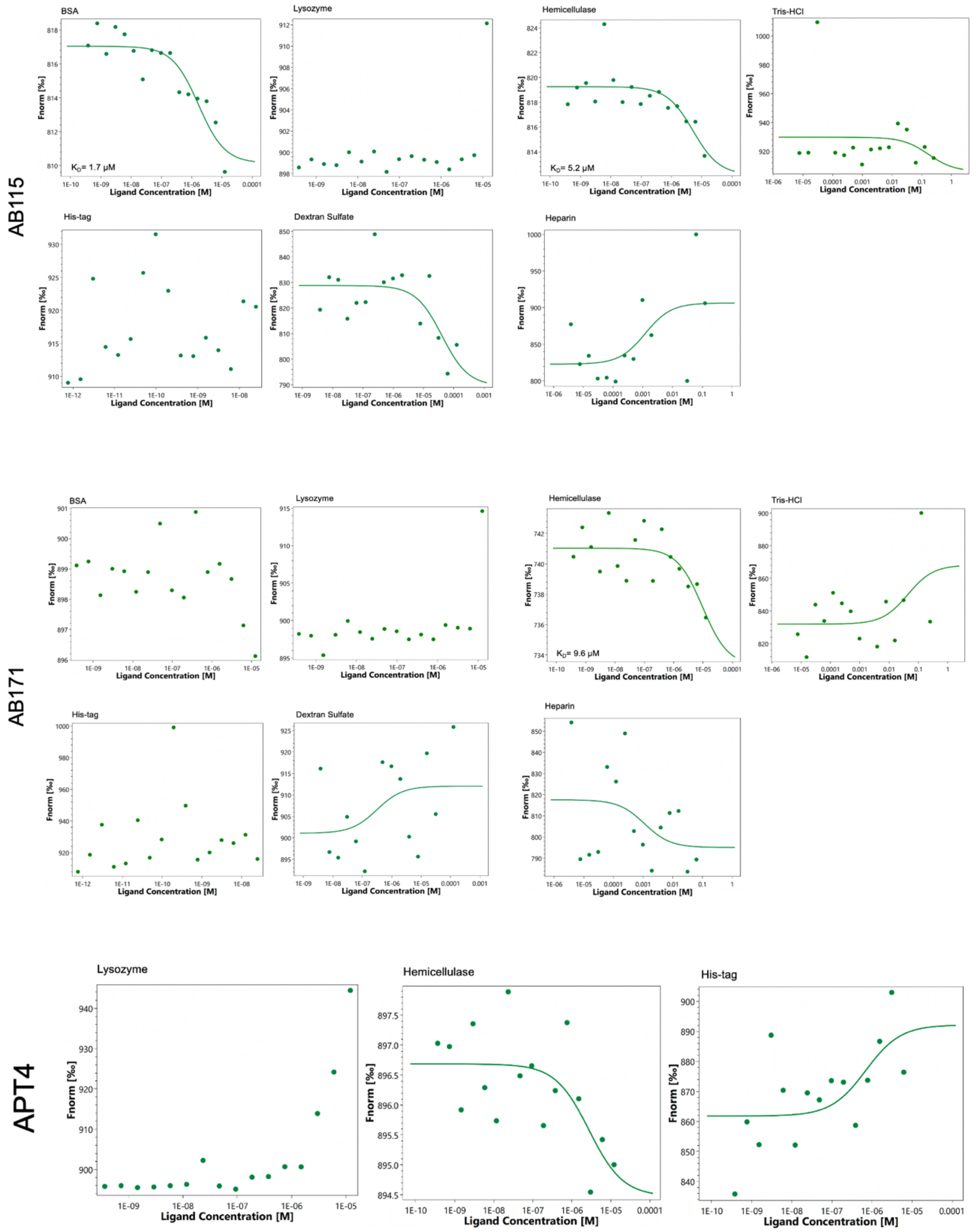

*Specificity test*

Binding specificity of selected candidates; ~5 nM of each candidate in presence of 12 µM to 0.*5* nM non-interest target.

BSA, lysozyme, hemicellulase and protease TEV (the latter in the presence of polyhistidine-tag, which was present in our ligand during candidate selection) were used in order to verify the specificity of the aptamers for PirAB. It was not possible to determine $K_D$ values for these "negative control proteins" (or the signal to noise ratio was not adequate), confirming specificity against the toxin. No curve obtained for lysozyme binding.
